# Molecular epidemiology of Colombian *Histoplasma capsulatum* isolates shows their polyphyletic behavior and point out raw chicken manure as one of the infections sources

**DOI:** 10.1101/449876

**Authors:** Luisa F. Gómez, Myrtha Arango, Juan G. McEwen, Oscar M. Gómez, Alejandra Zuluaga, Carlos A. Peláeze, Jose M. Acevedo, María L. Taylor, María del P. Jiménez

## Abstract

The thermally dimorphic fungus *Histoplasma capsulatum* is the causative agent of histoplasmosis, which is the most prevalent endemic mycosis in America. The replacement of organic matter in agro-ecosystems is necessary in the tropics, and the use of organic fertilizers has increased. Cases and outbreaks due to the presence of the fungus in these components have been reported. In Colombia, chicken manure is the most common raw material in the organic fertilizers production. In this work, we reached the isolation of the fungus from a chicken manure. Then, we were able to compare genetically 3 environmental isolates with 42 Colombian human clinical isolates. The genetic comparison showed the environmental isolates grouping together with the clinical isolates. This result suggests chicken manure as one of the infection source with *H. capsulatum*. Also, the phylogenetic analysis using another *H. capsulatum* isolates from databases showed that the Colombian isolates widely distributed in the relation tree. This result pointed out the great genetic diversity among *H. capsulatum* Colombian population.

## Introduction

Histoplasmosis is a disease caused by the thermally dimorphic fungus *Histoplasma capsulatum (H. capsulatum)* that has been documented in all continents except Antarctica. In America, it is highly endemic, particularly in the Ohio and Mississippi river valleys. Estimates based on intradermal skin tests indicate that approximately 90% of the people has been in contact with the fungus (1–4).

*H. capsulatum* infection occurs when a contaminated area is disturbed, which causes aerosolization and subsequent inhalation of infective hyphal fragments and microconidia. Once these particles are inhaled and reach alveoli, the body temperature stimulates a switch from mycelium to yeast form. Histoplasmosis disease development depends on host factors such as immune system and lung conditions as well as fungal factors such as virulence and the amount of inhaled infecting particles. This interaction gives rise to different clinical forms of the disease, ranging from an asymptomatic form to a severe disease that can be lethal (1–4).

*H. capsulatum* was first isolated from soil in 1945 by Emmons (5), who after observed an environmental association between the fungus and some animals such as bats and hens (6, 7). Later, other researchers, through the study of outbreaks, identified sources of infection and validated the relation between *H. capsulatum* with bats and birds manure (8–14). These studies made it possible to delineate populations at risk of histoplasmosis, since most cases have been described in ecotourists, speleologists, archaeologists, construction workers, poultry farmers, chicken manure collectors, organic fertilizers producers, farmers, and in general, all whose handle organic fertilizers. For the above, histoplasmosis has been defined as an occupational and recreational disease (12, 15–24).

In Colombia, Histoplasmosis report is not mandatory, nevertheless outbreaks and cases has been described. Some of the outbreaks have been related with the handle of organic fertilizers and its raw materials. Ordoñez et al. in 1997, made the compilation of 12 outbreaks occurred in the Colombian Andean Region, in 10 of those outbreaks the infection source was identified and in 2 the infection source were the handle of chicken manure contaminated with *H. capsulatum*. Then in 2002, Jimenez et al. described an outbreak comprised a family who get infected after fertilized a plant with a soil enriched with a *H. capsulatum* contaminated chicken manure. In Colombia, a total of 18 outbreaks of Histoplasmosis has been described, on those 415 people get exposed and 188 (45.3%) were identified as infected. The source of infection were mostly related with environmental sources especially caves visiting and chicken manure handling (16, 20).

In the tropical agro-ecosystems the reconstitution of the organic matter is needed, then organic fertilizers are widely used. In Colombia, the most common raw material in the production of organic fertilizers and amendments is chicken manure. Therefore, if *H. capsulatum* is found associated with chicken manure and also this is the principal raw material for organic fertilizers production in Colombia, then there are a lot of people exposed to the fungus.

In order to call the attention about the high frequency of *H.capsulatum* in tropical environments and make mandatory the histoplasmosis report and to design preventive measures addressed to protect the people who are mainly exposed like chicken manure collectors, organic fertilizers producers, or any people who handle these products, two steps are needed. First, we must to demonstrate the presence of the fungus in those substrates and second, we must to prove the relation between the isolates obtained from organic fertilizers or chicken manure with the isolates obtained from human clinical cases. In a previous work, we developed and applied a protocol based on the Hc100 nested PCR to search the genetic material of *H. capsulatum* in composted organic fertilizers, soils samples from caves and bird excretes. Then we detected *H. capsulatum* DNA in 10% of the tested samples (25). The present work was addressed to isolate *H. capsulatum* from Colombian environmental samples that tested positive by Hc100 nested PCR, in order to compare environmental isolates of the fungus with the Colombian clinical ones based on genetic differences by Multi-Locus Sequencing Typing (MLST).

Since the 1980’s, numerous studies have evaluated the genetic variation of *H. capsulatum* population using typing techniques such as Restriction Fragment Long Polymorphism (RFLP) (26–30), Variable Number of Tandem Repeats (VNTR) (31), MultiLocus Sequence Typing (MLST) (32) and Whole Genome Sequencing (WGS) (33). These studies have consistently shown a strong association between phylogenetic clusters and geographic origin. The currently accepted taxonomy indicates there are 8 distinct groups of *H. capsulatum*, based on MLST analysis of over 130 isolates with focus on four single copy genes including fatty acid desaturase (ole), β tubulin 1 (tub), ATP ribosylation factor (arf) and H antigen (H -anti) (34, 35). However, as with many microorganisms, taxonomic status remains dynamic with even more recent research arguing *H. capsulatum* should be broken up into 4 completely separate species: *H. mississippiense*, *H. ohiense*, *H. suramericanum* and *H. capsulatum sensu stricto* (Sepulveda et al. 2017).

The *H. capsulatum* Colombian population has been represented in previous typing studies by 16 Colombian clinical isolates. Interestingly, these isolates have shown marked differences among them relative to other *H. capsulatum* groups. The works by Kasuga et al. (1999, 2003), and Teixeira et al. (2016) show a high degree of variability among the Colombian isolates by MLST. The Kasuga’s work placed the Colombian isolates in Latin American clades (LAm) LAm A, LAm B and 2 solitary lineages. Similarly, the work achieved by Teixeira et al. (2016) placed the Colombian isolates in the phylogenetic species LAm B1, LAm A1 and LAm A2. Finally, Sepulveda et al. (2017), evaluated two additional Colombian clinical isolates, MV3 and MZ5, which they argue are so divergent that they should be reclassified into two different species: *H. capsulatum sensu stricto* and *H. suramericanum*, respectively. It is important to point out these two isolates are different from the 16 previously used in other molecular epidemiology or speciation works. Despite the fact that only a small number of isolates have been studied, the existing data indicate a great diversity among the *H. capsulatum* Colombian population (31–35).

Here we showed the presence of *H. capsulatum* in the Colombian environment and describe the genetic analysis of *H. capsulatum* Colombian population by MLST using both clinically relevant molecular markers as well as the sequences of genes used for diagnosis purposes including the partial sequence of 100 KDa protein (Hc100), M antigen (AgM) and two Sequences Characteristic Amplified Randomly (SCAR) (34–39).

## Materials and Methods

### Search for *H. capsulatum* Colombian isolates from environmental sources

#### Environmental samples collection

Samples were collected from organic fertilizers, which were the compost products from an organic matter source (i.e., food scraps, pruning material, straw, or sawdust) and a nitrogen source (i.e., excrement from animals, primarily poultry or other birds), soil from cave floors and bird and bat droppings. Between 500 and 1000 grams of each environmental sample was placed in a plastic zip lock bag and labeled with the date, geographical coordinates, and the type of sample (raw material, composted material, cave soil, bat excretes or chicken excretes). Samples were then sent to the Grupo de Micología Médica, School of Medicine, Universidad de Antioquia to be processed.

393 samples were collected between 2010 and 2017. Of these 393 samples, 273 (69.5%) were composted fertilizers and organic amendments. A total of 120 (30.5%) samples did not have composting treatment. In the last group, 21 (5.3%) samples were from caves floors and/or bat droppings and a total of 99 (25.2%) samples were from bird depositions or chicken manure.

#### DNA extraction from environmental samples using the “FastDNA SPIN Kit for Soil”

The FastDNA SPIN Kit For Soil^®^ (MP Biomedicals, Santa Ana, CA, USA) was used to extract DNA from environmental samples using the manufacturer’s instructions with some modifications. Briefly, the extraction was performed using the supernatant obtained from the suspension of 10 g of sample in 30 ml of saline solution containing 0.001% Tween 80 and 0.1% antibiotics (gentamycin and penicillin). The suspension was stirred vigorously for 1 min and allowed to settle for 20 min. This procedure was repeated twice. After the last stirring, the suspension was allowed to settle only until the largest particles were settled; then, 300 µl of supernatant was collected for DNA extraction. The other modification consisted of an increased contact time between the sample and kit reagents (25).

#### Nested Hc100-PCR assay for searching H. capsulatum in environmental samples

Two sets of specific primers targeting a fragment of the *Hcp100* gene were used (36, 37). The conditions of the assay for organic fertilizers were described by Gomez et al. (2018) (25). In the first reaction, 10µl of DNA was added to 50µl total reaction mix with a final concentration of 10 mM Tris - HCl (pH 8.3), 50 mM KCl, 2 mM MgCl2, 1U Taq polymerase (Thermo Scientific, Ref: EP0402. Waltham, M A, USA), 0.2 mM of each primer (HcI-HcII), and 0.2mm deoxynucleoside triphosphate (Thermo Scientific, Ref: R0181. Waltham, M A, USA). The mixture for the nested PCR was similar, except using 2μl of the product of the first PCR and 0.2mM of the inner primers (HcIII-HcIV). Temperatures and times for the first reaction, containing external primers HcI-HcII were one cycle at 94°C for 5 min; 35 cycles at 94°C for 30s, 66°C for 1 min, and 72°C for 1 min, and then a final cycle of 72°C for 5 min. For the second step, the reaction consisted of a cycle at 94°C for 5 min; 35 cycles at 94°C for 30s, 65°C for 30s, and 72°C for 1 min, and then a final extension cycle at 72°C for 5 min. Synthesis of the primers was performed by Integrated DNA Technologies (IDT, Coralville, IA, USA).

#### Agarose gel electrophoresis

Agarose gels (Amresco, Ref.: N605-500G, Solon, OH, USA) prepared at 1.5% in Tris-borate EDTA buffer (TBE) were used to visualize the amplification products of the Hc100 nested PCR. Electrophoresis was performed for 40 min at 80 V with 10 µl of the PCR product and 5 µl of GelRed Nucleic Acid Gel stain (Biotum. Ref.: 41003, Hayward, CA, USA) in each lane. The bands were visualized and documented in a UV transilluminator (Doc Gel ™, BioRad. Hercules, CA, USA).

#### H. capsulatum recovery in microbiological cultures from environmental samples that tested positive in the Hc100 nested PCR

The culture was performed using the supernatant obtained from the suspension of 10 g of sample in 30 ml of sterile saline solution 0.85% containing 0.001% Tween 80 (Sigma, ref. P4780. St. Louis, MO, USA), 100 pg/ml oxytetracycline (MK. Cali, Valle, COL), and 0.1% antibiotics (gentamycin and penicillin (MK. Cali, Valle, COL)). The suspension was stirred vigorously for 1 min and allowed to settle for 20 min. This procedure was repeated twice. After the last stirring, the suspension was allowed to sit until the largest particles were settled; then, the supernatant was collected and serial dilutions were performed 1:10, 1:100 and 1:1000, from each dilution were plated 200 µl in Mycosel agar (BBL^TM;^ ref. 211462. Franklin Lakes, NJ, USA) by duplicate. Cultures were incubated at room temperature for 2 months and plates were visually inspected on days 5, 10, 15, 30, 45 and 60 in order to look for growth of colonies with morphology that resemble *H. capsulatum.*

The colonies resembling *H. capsulatum* were identified as cottony colonies, raised, hard- edged, white, cream or light coffee and slow growth. Each candidate colony was subcultured on a fresh Mycosel agar and the microscopic examination was done with Lactophenol blue to observe the characteristics indicative of *H. capsulatum* such as thin septate hyaline hyphae, thin wall microconidia and tuberous macroconidia. Colonies with these features were transformed into the yeast form using the procedure described in Gomez et al. (2018) (25). Isolate identities were confirmed with sequencing of *hcp100* gene.

#### H. capsulatum recovery in mouse model from environmental samples that tested positive in the Hc100 nested PCR

##### Assays conducive to stablish the sensitivity of the mouse model

The efficacy of isolating *H. capsulatum* from the environmental sample was evaluated with organic fertilizers samples contaminated with 3000 CFU/ml in mycelial stage (Gomez et al. in preparation manuscript). 1:10, 1:100 and 1:1000 dilutions were prepared from the sample supernatant and from each dilution, 2 mice were inoculated in the peritoneum cavity with 500 µl. The animals maintenance is explained below.

#### Environmental samples preparation for mice inoculation

Hc100 nested PCR positive samples were prepared by suspending 10 g of sample in 100 ml of saline solution containing 0.1% antibiotics (gentamycin and penicillin). The suspension was stirred vigorously for 1 minute and allowed to settle for 20 minutes. This procedure was repeated twice. After the last stirring, the suspension was allowed to settle 40 minutes; then, the supernatant was collected for mice inoculation.

#### Animals’ maintenance and inoculation

Specific pathogen-free male Balb/C mice (6–8 week old, 18–22 g weight) were obtained from the Laboratory Animal Center at the Corporación para Investigaciones Biológicas (CIB, Medellin, Colombia). Mice were housed in a caging system (RAIR HD Super Mouse 750TM Racks system, Lab Products, Inc. Seaford, DE, USA) equipped with high efficiency particulate air (HEPA) filters with controlled room temperature at 20-24°C; and 12-h light/dark cycles, under sterile conditions and provided with sterilized food and water *ad libitum*.

Mice were inoculated in the peritoneum cavity with 500µl of the supernatant. After 17 days, mice were euthanized and the affected organs (spleen, liver and lungs) were extracted. The organs were macerated and cultured in BHI agar (Brain Hearth Infusion Agar, BBL ^TM^; ref. 211065. Franklin Lakes, NJ, USA) supplemented with 1% glucose (Sigma,Ref. G5400. St. Louis, MO, USA), 0.001% L-cysteine (Sigma, ref. C - 7755. St. Louis, MO, USA) and 5% anticoagulated blood, then cultures were incubated at 37°C in 5% CO_2_ atmosphere for two weeks and at room temperature in Mycosel agar for 2 months, in order to look for *H. capsulatum* compatible yeast colonies.

#### Ethics statement

This study was performed according to recommendations of European Union, Canadian Council on Animal Care, and Colombian regulations (Law 84/1989, Resolution No. 8430/1993). The protocol was approved by the Ethics’ Research Committee at the CIB (Comité de ética, electronic consultation on July 24th, 2014).

### Collection of Colombian *H. capsulatum* clinical isolates

#### H. capsulatum culture

42 human clinical isolates of *H. capsulatum* were obtained from the CIB (CIB, Medellin, Colombia). All isolates were transformed into the yeast form following the procedure were described in Gomez et al. (2018) (25).

#### DNA extraction from H. capsulatum isolates in yeast phase

The phenol-chloroform-isoamyl alcohol method was used for DNA extraction from the *H. capsulatum* isolates in yeast phase (40). The DNA concentration and quality were evaluated with a NanoDrop ND1000 spectrometer (Thermo Scientific, Wilmington, DE, USA) and 1% agarose gel electrophoresis, respectively

### Molecular epidemiology comparison among *H. capsulatum* population

#### MLST technique

The characteristics of the technique to compare the isolates of *H. capsulatum* were described by Kasuga, Taylor and White (1999) (34). The primers were synthesized by Integrated DNA Technologies (IDT, Coralville, IA, USA). Table S2 describes the characteristics of each primer.

The standardized conditions for each MLST PCR assay and primers used are described on Table S2.

#### Other selected and evaluated H. capsulatum genome regions to compare isolates

Given the difficulties associated with obtaining isolates from environmental samples, diagnostic protocols originally designed for clinical samples were standardized and applied in environmental samples to obtain sequences directly from the samples to enable comparison between clinical and environmental *H. capsulatum* isolates. To fulfil this goal only the first round of the Hc100 Nested PCR and antigen M nested PCR assays were done because the amplification product is bigger. Additionally, two PCR assays targeting two Sequences Characteristic Amplified Randomly (SCAR) in the *H. capsulatum* genome were also used. Table S2 describes the characteristics of each primer and PCR reaction.

#### Sequences analyses

Bidirectional sequencing of the PCR amplification products were performed using the chain termination method. The method used ABI 3730XL DNA sequencing technology with quality criteria QV20 (Macrogen Inc., Geumcheon-gu, Seoul, Korea). The obtained sequences were edited manually based on the chromatograms. The mold and complementary sequences were aligned and the consensus sequence was obtained using Geneious 11.0.2. Software. BLAST (http://blast.ncbi.nlm.nih.gov/Blast.cgi) was used to verify that the sequenced PCR products belonged to *H. capsulatum.* It was obtaining the information of the 80 sequences from TreeBASE #1063, the matrix was separated gene by gene according to the length and the order reported by the author (34). Additionally, the sequences of 85 isolates were downloaded from NCBI using the access codes by the Batch entrez program. Additionally, the sequences were aligned and concatenated using Geneious 11.0.2. Software, and the distance trees were constructed using the Maximum Likelihood and 10000 UltrafastBootstrap methods using IqTree 1.4.4. Software. The tree obtained was visualized using FigTree v.1.4.3.

## Results

### Detection of *H. capsulatum* in environmental samples by Hc100 nested PCR

Out of the 393 environmental samples, a total of 39 (9.9%) were positive by Hc100 nested PCR. Of these positive samples, 21 (5.3%) were composted fertilizers and organic amendments, 17 samples (4.3%) were from bird depositions or poultry manure and one (0.3%) sample was from a cave floor/bat droppings. Interestingly, we found high positivity rates of samples without composting treatment 18/120 (15%) compared with the samples that were previously composted 21/273 (7.7%).

### Recovery of *H. capsulatum* in microbiological cultures from Hc100 nested PCR positive environmental samples

The 39 positive samples by the Hc100 nested PCR were seeded into Mycosel culture medium. However, only one sample taken from non-composted chicken manure produced viable *H. capsulatum* cultures. From this single sample, 3 *H. capsulatum* colonies were obtained and used as different individuals for molecular analyses. Interestingly, while 2 of these isolates were morphologically similar, the third was distinct and did not fully switch to the yeast form. Abundant growth of bacteria and molds, such as *Penicillium* spp*, Aspergillus* spp*, Trichoderma* spp and *Geotrichum* spp were also observed in the *H. capsulatum* cultures.

### Recovery of *H. capsulatum* in mouse model from Hc100 nested PCR positive environmental samples

In attempt to obtain more isolates, the 39 positive samples from the Hc100 nested PCR were also used to inoculate mice. 5 to 8 mice were used to inoculate each positive sample, around 9% of the mice died in the first 3 days after inoculation. Despite the use of broad-spectrum antibiotics such as oxytetracycline in the sample preparation, the administration of antibiotics to the mice before and after environmental samples inoculation and the rigorous observation of the mice tissue cultures in Mycosel and BHI, we were not able to obtain environmental isolates of *H. capsulatum*, although as it was described before *H.capsulatum* was recovered from mice inoculated with environmental samples artificially contaminated with 3000 CFU/ml of this fungus.

### MLST analysis of Colombian *H. capsulatum* populations

The conditions of each PCR protocol were standardized (Table S2). Then the sensitivity, specificity and the ability to detect *H. capsulatum* DNA from environmental samples could be evaluated (Figure 1). All protocols, including both typing and diagnostic methods, were tested to detect *H. capsulatum* from environmental samples. Although some of them showed a good limit of detection (Fig 1, Lanes 1-7) and specificity (Fig 1, Lanes 8-13) with spiked samples with *H. capsulatum* DNA, none of these methods produced an amplicon from the 39 samples shown to be positive with the Hc100 nested PCR.

**Figure 1.**
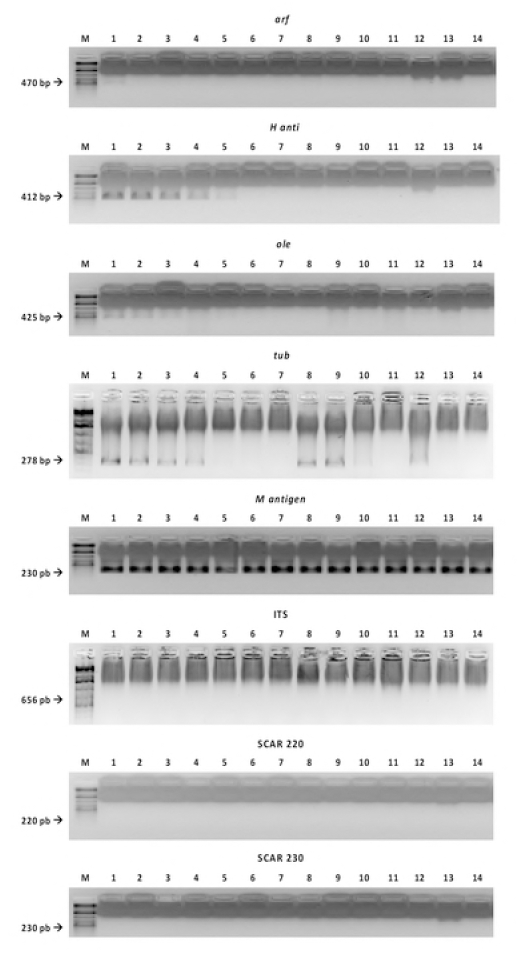
Detection limit, specificity and the ability to detect *H. capsulatum* DNA in environmental samples with the MLST and diagnostic PCR reactions. Line M: marker, line 1: 2 ng/µl, line 2: 1 ng/µl, line 3: 500 pg/µl line 4: 200 pg/µl, line 5: 100 pg/µl, line 6: 50 pg/µ1, line 7: 20 pg/µl line 8: *Paracoccidioides brasiliensis* sensu stricto (Pb 18), line 9: *Paracoccidioides* spp (Pb 339), line 10: *Paracoccidioides restrepiensis* (PB 60855), line 11: *Coccidioides* spp, line 12:

The standardized PCR protocols were used to obtain the sequences from all 45 Colombian isolates (42 human clinical and 3 environmental isolates) (Table S1), after the PCR products were sequenced, the consensus sequence was obtained. The length for each locus was as follows: *arf* 478 bp, *anti H* 413 bp, *ole* 428 bp, *tub* 263 bp, *100kda* 209 bp, *anti M* 276 bp, scar220 208 bp, scar230 262 bp, ITS 605 bp. The characteristics of each gene are shown in the Table S2.

Subsequently, the first matrix was constructed using the sequences of the genes in the following order: *arf*, *antiH*, *tub1* and *ole*. The matrix included 225 sequences, had a length of 1582 bp and was partitioned by the type of sequence itself: CDS, exon or ITS. The evolutionary model was established as K2+G4. 17 phylogenetic groups were identified with similar distribution as reported by Kasuga, 2003 and Teixeira, 2016 (Figure 2).

**Figure 2.**
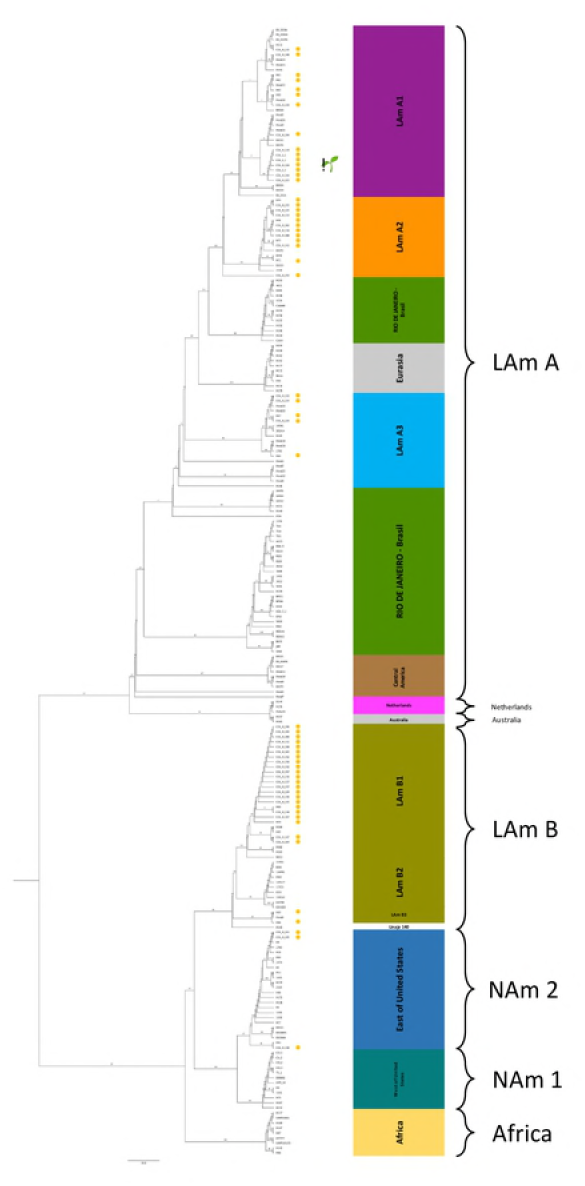
Genetic comparison *of H.capsulatum* Colombian isolates. The groups obtained, were separated when bootstraps were over 70%. The contrast between our results and the Teixeira et al. (2016) (middle list) and the Kasuga et

Additionally, three more matrices were built that only included the Colombian isolates, given that only from those were obtained the sequences corresponding to Hcp100 gene, M antigen gene, SCAR 220, SCAR 230 and the ITS region. In this analysis, the first matrix included *arf*, *antiH*, *tub1* and *ole*. The second matrix included the sequences obtained using the PCR products of the diagnosis genes target, those sequences were aligned in the order as shown above and a third matrix included all nine sequences. In all matrices, the environmental isolates were grouped with the clinical isolates. When the 3 different matrices were analyzed, the trees had the same topology, this suggest that the classical genes are adequate to achieve the genetic comparison inside the *H. capsulatum* population (Figure 3). According to the genes characteristics for MLST, the diagnostic genes have less number of informative sites per gene than the classical targets (Table 1).

**Figure 3.**
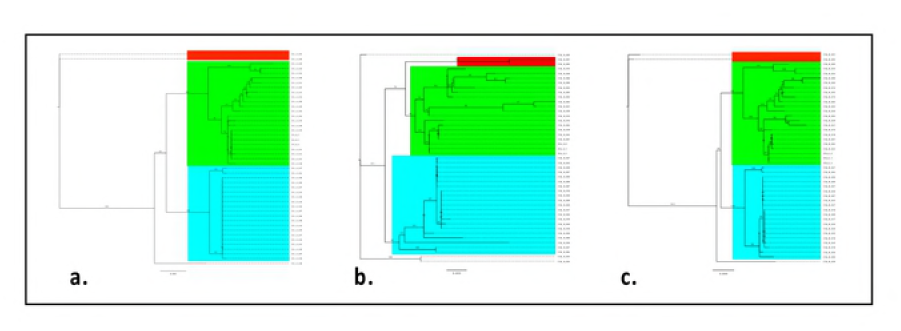
The phylogenetic analysis of the *H.capsuhtum* Colombian population. All trees showed the same topology although there were built with different group of genes in the order as follow, a. *arf, antiH, tubi* and *ole.* b. HcplOO gene, M antigen gene, SCAR 220, SCAR 230 and the ITS region and c. Consensus tree of all 9 sequences *arf, antiìi, tubi, ole,* HcplOO gene, M antigen gene, SCAR 220, SCAR 230 and the ITS region.

**Table 1.**
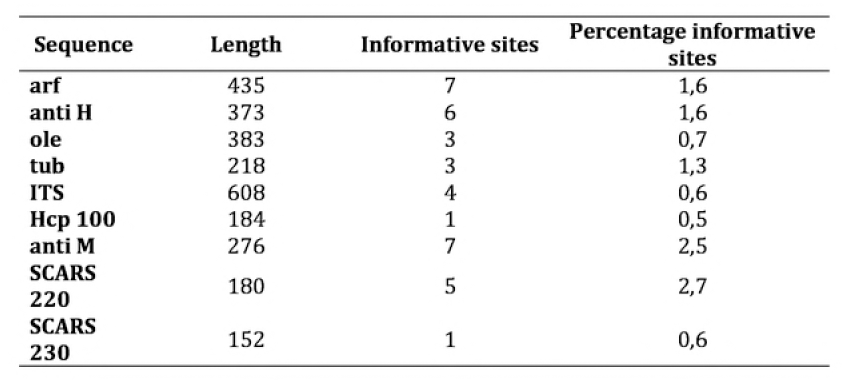
List the sequence name, the length in the present analysis and the number and percentage of the informative sites.

## Discussion

Histoplasmosis is the most prevalent endemic mycoses in America. *H. capsulatum*, the causative agent, is a thermally dimorphic fungus associated with bat/cave environments, bird droppings, and other diverse environmental sources such as soil and compost.

People exposed to *H. capsulatum* can be either asymptomatic or experience life threatening histoplasmosis depending on a combination of risk factors (4, 41). A better understanding of the genetic diversity of environmental and clinical isolates *H. capsulatum* in context of geographical location can help us better understand the risk factors for developing histoplasmosis. In this study, we showed how *H. capsulatum* strains in Colombia show a particularly high degree of variation compared to other geographic regions through the phylogenetic analysis of over 3 environmental and 42 human clinical isolates.

Using Hc100 nested PCR, we were able to detect *H. capsulatum* in 39 (9.9%) of 393 of environmental samples studied. The fact that of the positive samples, 21.9% were found in non-composted samples compared to only 7.7% of composted samples, suggests that a well-done composting process could reduce the risk of exposure to *H. capsulatum* when manipulating the organic fertilizer final product. This is because composting process is a spontaneous decomposition of organic matter, mainly aerobic, in which bacteria and fungi mainly participate. The organic matter is transformed in a free of toxins and pathogenic microorganisms fertilizer due to the action of the saprophyte microorganisms. First, they are better competitors for space and nutrients and second due to their metabolic activity, which generates an increase in the temperature. These two actions reduce the pathogenic microorganisms population (42-44). However, the composting process at the beginning, continuing being a high risk infection source, since we achieve the isolation of *H. capsulatum* from a raw/not composted chicken manure, that happens to be the most used raw material in organic fertilizers in Colombia.

For the isolation of *H. capsulatum* from environmental samples, the mouse model has been considered the golden standard technique, but, in this study, we could not isolate the fungus using it. This procedure is time-consuming, expensive, requires training to handle the mice and based on previous studies it has a low success rates between 0-50% (5, 6, 8, 19, 20, 45-51). Furthermore, we found the high microbial background in our environmental samples led to sepsis and subsequent death in many of the mice, further emphasizing the high cost of such experiments that ultimately failed to recover the *H.capsulatum* isolates. Moreover, the search for *H. capsulatum* using the mouse model requires the fungus to overcome the saprophytic microorganisms, evade the immune system and establish an infection in the mammal host before it can be isolated from affected tissues (5, 6, 8, 19, 20, 45-51). One challenge associated with this method is that if the fungus has been in the environment for long time without the exposure to a mammal host, the genes associated with pathogenicity may not be activated. For example, when the α-(1–3)-glucan cell wall content has been studied in *H. capsulatum,* the fungus has been separated into chemotypes I and II; this classification has been related also with its virulence. Most *H. capsulatum* isolates described are chemotype II, but can lose virulence spontaneously by successive passaging in culture by loss of α-(1–3)-glucan in their cell wall (52-54). Nowadays, it is possible to compare isolates using the proteome analysis. This approach could be used to elucidate genes related to pathogenicity by comparing gene activity in environmental and clinical isolates (55-57). If environmental isolates have not activated the virulence genes like the clinical isolates, then the use of the mouse model for the isolation of *H. capsulatum* from environmental samples would need to be reconsidered to be the gold standard technique .

In contrast, we were able to isolate *H. capsulatum* by the direct culture of the environmental samples. In cases with high microbial background such as chicken manure samples, we were able to limit contamination by adding antibiotics to the culture media. Even though this method also had a low success rate, the recovery of just a few isolates proved to be very valuable for our study. From our analysis of the 3 isolates from the non-composted chicken manure sample, we were able demonstrate viable fungus was present in the raw materials. This has implications for worker safety in industrial contexts, and its necessary take actions regarding protective measures like wearing high efficient respirators, gloves, boots, long sleeve shirts and showering at the end of the day. Following these few recommendations is the best way to avoid the occurrence of occupational outbreaks of histoplasmosis (58). In order to promote the knowledge about histoplasmosis and how to avoid the infection with *H. capsulatum,* we wrote a booklet to teach people about the disease, the signs and symptoms, whom is usually expose and how to protect themselves (59).

Given the difficulties associated with obtaining isolates from environmental samples, we probed diagnostic protocols probed, to obtain sequences directly from the environmental samples to enable comparison between clinical and environmental *H. capsulatum* isolates. However, none of these methods produced an amplicon from the 39 samples shown to be positive with the Hc100 nested PCR. Also, given that the trees obtained with the 3 different had the same topology, the classical genes proposed by Kasuga et al. are adequate to achieve the genetic comparison inside the *H. capsulatum* population (34, 35, 37–39, 60).

Then, Our successful recovery of three *H. capsulatum* isolates from the chicken manure sample enabled us to genetically compare these strains to 42 human clinical isolates from Colombia, as well as other isolates from around the world by the classical MLST (34, 35). Our MLST analysis revealed that the isolates from chicken manure are most closely related to the clinical strains in Colombia. This finding strongly implicates chicken manure as a source for exposure and infection by *H. capsulatum*. Previous studies of histoplasmosis outbreaks have used similar approaches and also been able to establish associations between environmental and clinical *H. capsulatum* isolates (26, 32, 35).

Our work confirming the relatively high diversity of Colombian isolates compared to populations in other geographical regions (32, 33, 35). Although it is not yet certain why there is such a high diversity, it is likely related to Colombia’s ecological and geographical characteristics. Colombia has a unique location that serves a crossing point for migratory birds and bats, which can carry and seed the fungus. This mechanism of dispersal has been demonstrated by studies of the bat *Tadarida brasiliensis*, whose range correlates well with the epidemiological map of histoplasmosis. This mechanism may explain the presence of

Colombian isolates in the clades that traditionally include isolates from other countries the bat visits in its migratory route like the United States and Brazil. Furthermore, *H. capsulatum* has also been isolated from tissues and excretes from both birds and bats (51, 61–66). Also, since tropical soils tend to be nutritionally poor, organic fertilizers made on chicken manure sources are commonly used, which creates increased habitat for *H. capsulatum* that overlaps with human activity. Our data implicates a mechanism of clinical infection originating from non-composted chicken manure sources. It is therefore likely that the risk of histoplasmosis can be reduced through complete composting of chicken manure before extensive handling by people. Risk may also be reduced by, the use of personal protection when exposed to risky activities like the visit to caves, the demolition or cleaning of old buildings or the manipulation of excreta of birds and bats and the inclusion of the *H. capsulatum* search during the composting process. Following these recommendations, cases and outbreaks could be prevented.

## Acknowledgments

This study was supported by Comité para el Desarrollo de la Investigación (CODI) of Universidad de Antioquia, project code 2014-1010 and Departamento Administrativo de Ciencia, Tecnología e Innovación (Colciencias), Bogotá, Colombia, National Doctorate Program announcement 647 of 2014 to LFG. The funders had no role in study design, data collection and analysis, decision to publish or preparation of the manuscript. The authors would like to acknowledge to the members of the participant research groups for feedback and improvement suggestions. We thank to David Sexton for English editing of the manuscript.

## Data Accessibility

Genbank accession numbers ADP ribosylation factor (arf) MH338036 to MH338080, H antigen precursor (*anti-H*) MH122839 to MH122883, Delta-9 fatty acid desaturase (*ole1*) MH338126 to MH338170, Alpha-tubulin (*tub1*) MH338081 to MH338125, Hcp100 gene, 100 kDa protein MH122794 to MH122838, internal transcribed spacer (ITS) MH339542 to MH339586, SCAR markers 220 and 230 MH348521 to MH348610.

